# Cell morphology deep learning guides discovery of mitochondrial host defense against influenza

**DOI:** 10.64898/2026.02.25.707858

**Authors:** Olha Shkel, Hyeonsu Kim, Md. Mamunul Haque, Yevheniia Kharkivska, Kyung Tae Hong, Sun-Hak Lee, Yun Kyung Kim, Chang-Seon Song, Woo Youn Kim, Jun-Seok Lee

## Abstract

Influenza A virus exploits host cellular machinery across subcellular compartments, yet the organelle-level changes that distinguish infected from uninfected cells and the molecular players driving them remain poorly defined. Here, we combine organelle image-based deep learning with proximity labeling chemoproteomics to address this gap. A convolutional neural network identified mitochondrial morphology as the strongest single-cell predictor of infection status (precision = 84.9%). Proximity labeling profiling of mitochondrial matrix proteome revealed 99 proteins with significantly altered upon infection, of which five (CH60, ETHE1, LONM, MPPB, and SQOR) were validated as host restriction factors whose depletion elevated interferon-β expression, enhanced viral RNA accumulation, or increased progeny virus production. Notably, two of these factors, ETHE1 and SQOR, operate within the mitochondrial hydrogen sulfide oxidation pathway, and pharmacological scavenging of H_2_S by hydroxocobalamin dose-dependently reversed their knockdown phenotypes, directly linking mitochondrial sulfide metabolism to antiviral defense against influenza.

## Introduction

Influenza A virus (IAV) continues to pose a significant threat to global public health, contributing to seasonal epidemics and periodic pandemics with significant morbidity and mortality (*1, 2*). The virus extensively exploits host cellular machinery throughout its replication cycle, orchestrating processes across subcellular compartments (*3–5*). Infected cells often undergo substantial morphological and functional remodeling, especially at the level of organelles such as the mitochondria and endoplasmic reticulum (ER). These alterations can impact antiviral signaling, metabolism, and protein trafficking (*6–8*).

Detecting such infection-induced cellular changes at the single-cell level has been a long-standing challenge. Chemical biology approaches have demonstrated that organelle-targeted fluorescent probes can capture early infection events with single-cell resolution. For example, we previously demonstrated that sulfinate-based superoxide probes enabled cell line-dependent discrimination of avian influenza virus infection through intracellular reactive oxygen species dynamics (*9*), and ER-targeting fluorophores detected early infection by monitoring organelle-specific stress signatures (*10*). Although these chemosensor-based strategies offer organelle-level specificity, they inherently depend on the design and synthesis of specialized molecular probes, which may limit their accessibility to the broader research community.

In parallel, advances in deep learning have opened a fundamentally different path toward single-cell phenotypic analysis. Convolutional neural networks (CNNs) can now predict subcellular fluorescence patterns directly from transmitted-light microscopy (*11*), segment and track organelle morphology in live cells (*12*), and localize organelles in label-free images by learning cell context (*13*). In virology, machine learning has been used to identify Zika virus-infected cells from changes in ER morphology (*14*) and to reveal subcellular remodeling caused by SARS-CoV-2 (*15*). These developments raise the possibility that infection-induced morphological fingerprints—phenotypic signatures potentially imperceptible to human observers—could be captured by image-based deep learning without requiring any exogenous probe.

Here, we combined image-based deep learning with spatially resolved proximity labeling proteomics to discover mitochondrial host factors that modulate influenza A virus infection. We first trained a CNN classifier on fluorescence microscopy images of individual organelles and whole cells, and found that mitochondrial morphology provided the strongest discriminatory signal for distinguishing infected from non-infected cells, establishing mitochondria as a primary site of infection-associated structural remodeling. This observation directed our subsequent molecular investigation toward the mitochondrial interior—a compartment that, despite the well-characterized role of outer membrane MAVS in innate antiviral signaling (*16*), remains largely unexplored in the context of influenza infection (*17–19*). To identify the molecular players within this compartment, we deployed APEX2-based proximity labeling targeted to the mitochondrial matrix, which enables compartment-specific protein mapping that conventional whole-cell proteomics or biochemical fractionation cannot achieve (*20, 21*). Quantitative mass spectrometry revealed 99 mitochondrial proteins with significantly altered abundance upon infection. Functional validation through siRNA-mediated knockdown demonstrated that five mitochondrial matrix and inner membrane proteins modulate the innate immune response and restrict viral replication.

## Results

### Deep learning-based infection prediction

We reasoned that if influenza A virus remodels host cell organelles, then morphological changes visible by microscopy might be sufficient to classify infection status without relying on viral marker immunostaining or specialized molecular probes such as superoxide or ER stress sensors (*9, 10*). To investigate how influenza A virus alters cellular ultrastructure, we collected single-cell microscopy images at both whole-cell level (bright-field) and organelle level, capturing nuclei, mitochondria, and endoplasmic reticulum (ER). Infection status was independently assigned by immunofluorescence against the influenza Matrix 1 protein: cells exceeding a defined intensity threshold were labeled infected, and all others as non-infected. We then trained a convolutional neural network (CNN) to predict infection status solely from these host cell images (Fig. 1A). The CNN successfully discriminated infected from non-infected cells across all image types, but performance varied markedly by organelle (Fig. 1B). Evaluated by area under the receiver operating characteristic curve (AUROC), mitochondrial images consistently outperformed all other inputs: AUROC = 0.854 for mitochondria, compared with 0.789 (nuclei), 0.723 (bright field), and 0.671 (ER) (Fig. 1C, Supplementary Table 1). This ranking result held even after down-sampling to match the smallest dataset size, with mitochondria-based classification retaining an AUROC above 0.8 and achieving the highest accuracy (79.7%) and precision (84.9%) on the independent test set. These results support mitochondrial morphology as the most informative visual signature of influenza infection at the single-cell level.

**Fig. 1.**
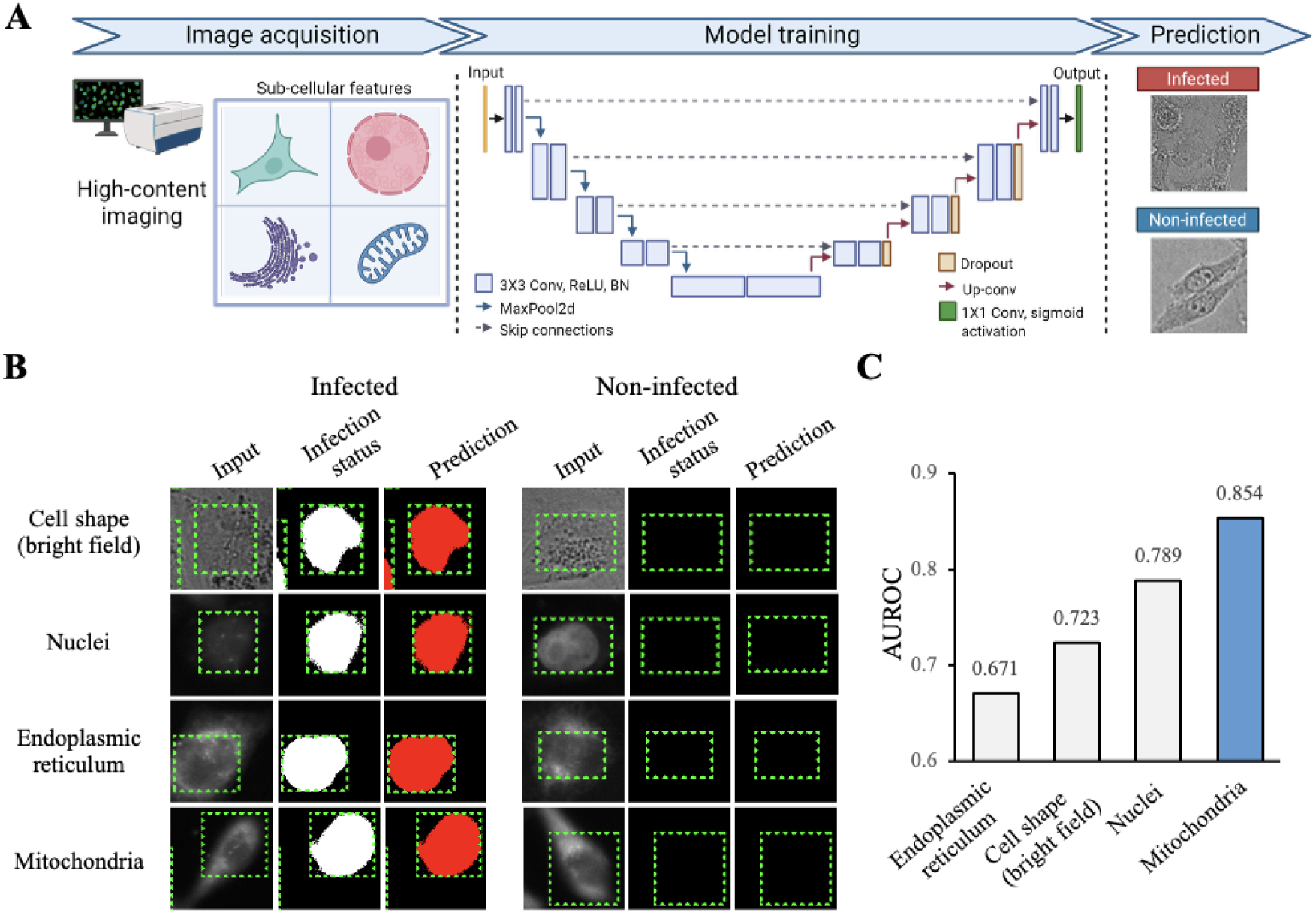
Deep learning classification of influenza infection from organelle morphology images. **(A)** Workflow of CNN model training. Bright-field, nuclear, endoplasmic reticulum, and mitochondrial images of influenza-infected and non-infected cells were acquired, processed into single-cell datasets, and used to train a convolutional neural network with an encoder-decoder architecture. The final dataset comprised 4,445 mitochondrial, 1,771 endoplasmic reticulum, 10,870 nuclear, and 10,870 bright-field images. The network employed ReLU activation in intermediate layers and a sigmoid output for binary classification, enabling the model to distinguish infected from non-infected cells. **(B)** Example of infection prediction. Left: model input image; center: infection status (white indicates infected); right: model prediction (red denotes cells predicted as infected). **(C)** Model evaluation. AUROC values for models trained on different image types are shown. Mitochondrial images provided the highest discriminatory power compared to cell shape (bright-field), nuclear, and endoplasmic reticulum inputs

### Proximity labeling chemoproteomics for mitochondria proteome profiling

Having identified mitochondrial morphology as the strongest imaging predictor of infection, we next sought to characterize the molecular changes occurring within this compartment. We employed proximity labeling to map the mitochondrial matrix proteome with spatial precision. APEX2, an engineered monomeric ascorbate peroxidase, generates short-lived biotin-phenoxyl radicals upon treatment with biotin-phenol and H_2_O_2_ that covalently tag nearby proteins (*20*). We targeted APEX2 to the mitochondrial matrix (Mito-APEX2) and confirmed its correct localization by colocalization with the mitochondrial marker TOM20 (Extended Data Fig. 1a): Mito-APEX2 and TOM20 signals showed strong spatial overlap, with a mean Pearson’s correlation coefficient of 0.790 ± 0.041 and Manders’ overlap coefficients of 0.877 ± 0.059 (M1, Mito-APEX2 overlapping TOM20) and 0.745 ± 0.102 (M2, TOM20 overlapping Mito-APEX2) (JaCoP, ImageJ, Supplementary Table 2). Treatment with 1 mM biotin-phenol for 1 hour followed by 1 minute of H□O exposure effectively labeled proteins in the APEX2-proximal region (Fig. 2A), and western blot analysis confirmed that Mito-APEX2 expression and labeling efficiency remained stable between infected and mock-infected conditions (Suppl. Fig. 2a, 2b).

**Fig. 2.**
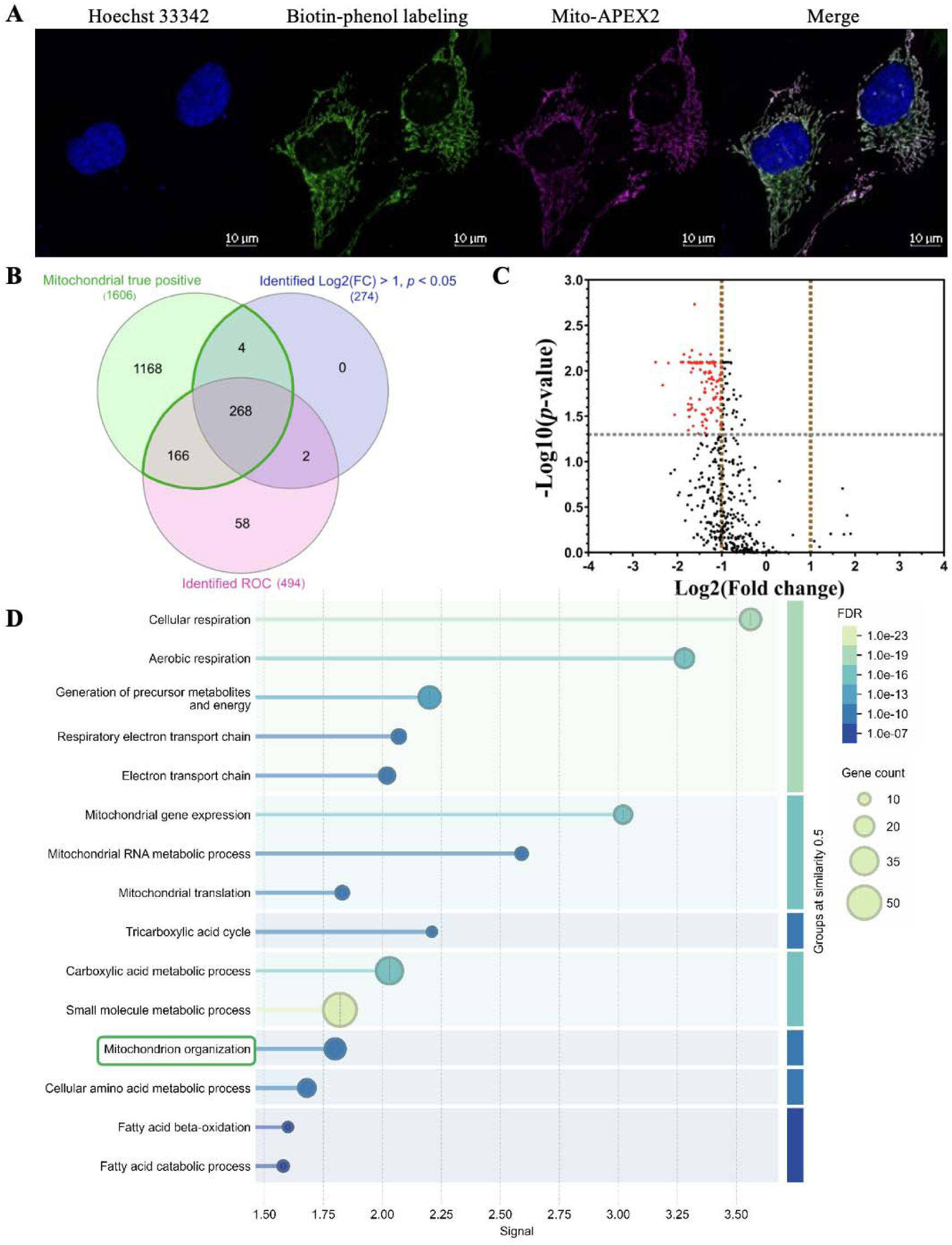
Proximity-labeling based mitochondria proteome profiling. **(A)** Confocal fluorescence images of HeLa cells expressing APEX2 targeted to mitochondria. Cells were treated with 1 mM biotin-phenol for 1 hour, followed by 1 mM H□O□ for 1 minute. Biotinylated proteins were visualized using Alexa Fluor 488 streptavidin conjugate, while expressed APEX2 was detected using an anti-V5 tag antibody. **(B)** Two sets of identified mitochondrial proteins were generated: (i) proteins with Log[(Fold change) > 1 and p < 0.05, and (ii) proteins selected based on ROC analysis. Each set was compared against a reference list of known mitochondrial proteins (‘Mitochondrial true positive’). The overlaps were visualized in a Venn diagram. Proteins falling within the intersection of each dataset and the true positive list were considered high-confidence mitochondrial proteins and were retained for further analysis (highlighted by the green enclosure). **(C)** Volcano plot of identified mitochondrial proteins. Log2(Fold change) ratios and -Log10(p-values) for APEX2-infected versus APEX2 mock-infected samples were plotted for each protein. Ninety-nine proteins with significant differential abundance ratios were identified (red). The threshold values for Log2(Fold change) were 1 and -1; the threshold for –Log10(p-value) was 1.3. **(D)** Enrichment analysis of differentially abundant proteins. The dot plot represents the top 15 enriched terms in the Biological Process (GO) category (full list is available in Supplementary Table 3). The terms are ranked based on enrichment signal and grouped by similarity (0.5). The term ‘Mitochondrion organization’ is highlighted in green.

To dissect infection-induced proteomic changes within the mitochondrial matrix, we designed a three-condition experiment comparing: (1) mock-transfected, mock-infected cells (Control), (2) APEX2-transfected, mock-infected cells (APEX2), and (3) APEX2- transfected, IAV-infected cells (APEX2 Infected), all subjected to identical follow-up protocols for biotinylations. The Control condition captures background biotinylation, the APEX2 condition defines the baseline mitochondrial matrix proteome, and the APEX2 Infected condition reveals infection-driven changes. This stepwise logic allows us to distinguish APEX2-specific labeling from nonspecific background and isolate the effect of infection. Each condition was processed in triplicate, the biotinylated protein were enriched on streptavidin beads, digested with trypsin, and labeled with TMT10plex™ isobaric tags for multiplexed quantitative proteomic analysis. After correction for isotopic impurities and normalization across TMT channels, we identified 1,486 proteins (unique peptides ≥ 2; Supplementary Table 3).

We next established a high-confidence inventory of mitochondrial matrix proteins captured by proximity labeling. Comparing protein abundance in mock-infected APEX2-labeled samples against non-labeled controls identified 274 significantly enriched proteins (Log (fold change) > 1, *p* < 0.05). Because simple fold-change thresholding can miss genuinely labeled proteins with moderate enrichment, we complemented this approach with a receiver operating characteristic (ROC) curve-based classification as previously reported: proteins were ranked by their TMT intensity ratio and evaluated against curated true-positive and false-positive mitochondrial reference sets with optimal cutoff chosen to maximize sensitivity while minimizing false discovery (*22*). This second method yielded 494 candidate proteins. Intersecting the enrichment-based and ROC-derived datasets with a reference list of 1,606 known mitochondrial proteins compiled from multiple databases produced 438 high-confidence mitochondrial proteins, which was used as the core dataset for all subsequent analyses (Fig. 2B, Supplementary Table 3). Of these 438 proteins, 99 showed significantly reduced abundance in infected cells relative to mock-infected controls (Log (fold change) ≤ –1, p < 0.05) (Fig. 2C), indicating a widespread yet selective remodeling of the mitochondrial proteome upon influenza A virus infection. Protein-protein interaction analysis (STRING v12.0) (*23*) revealed that these 99 proteins form a highly interconnected network (208 edges, average node degree of 4.2, clustering coefficient 0.572) enriched in core mitochondrial processes: cellular respiration (22 proteins, FDR = 2.00 x 10^-19^), mitochondrial gene expression (17 proteins, FDR = 4.59 x 10^-15^), and mitochondrial RNA metabolic process (10 proteins, FDR = 1.13 x 10^-10^) (Fig. 2D, Supplementary Table 4). Notably, 21 of the 99 proteins were annotated under ‘mitochondrion organization’ (FDR = 1.36 x 10^-11^), which is a term encompassing morphology, dynamics, genome replication, and biogenesis (Suppl. Fig. 3). This result directly resonates with the mitochondrial morphological changes that our CNN has identified as the strongest imaging signature of infection. These functionally interconnected clusters (Extended Data Fig. 2) suggest that influenza infection does not deplete mitochondrial proteins but instead targets coherent functional modules essential for organelle integrity and energy homeostasis.

### Validation of infection-induced mitochondrial proteome remodeling

The coordinated downregulation of 99 mitochondrial proteins from proximity labeling upon infection raised an important question: whether the observed changes reflect selective remodeling of the matrix proteome or a general reduction in mitochondrial content. Distinguishing between these two scenarios is essential as mitophagy, the selective autophagic degradation of mitochondria, has been reported during influenza replication stress and could account for a global decline in mitochondrial protein levels (*24, 25*). To address this, we examined established markers of mitophagy and mitochondrial mass in infected and mock-infected cells (Fig. 3A). PINK1, a key mediator of mitophagy initiation (*26*), was detected exclusively in its truncated form under both conditions, with no accumulation of the full-length species indicative of active mitophagy. Levels of TOM20 and VDAC, outer membrane proteins widely used as indicators of mitochondrial mass, remained unchanged upon infection. These data argue against mitophagy-driven organelle loss and instead indicate that influenza virus infection induces a selective reprogramming of the mitochondrial matrix proteome independent of changes in overall organelle content. On the basis of these findings, we selected nine downregulated proteins from proximity labeling for individual validation and functional characterization, prioritizing candidates annotated under the Gene Ontology term ‘mitochondrial organization’, which is the biological process most directly linked to the morphological remodeling detected by our analysis (Table 1, Suppl. Fig. 3). Six of these proteins (CH60, GRP75, LONM, MPPB, OXA1L, and TIM44) are involved in essential matrix functions including protein import, folding, proteolytic processing, and membrane insertion. LETM1 was included for its established role in cristae morphology and respiratory chain assembly, and ETHE1 and SQOR for their function in mitochondrial sulfide metabolism, which is a pathway required for cytochrome c oxidase activity and cellular homeostasis. Although these candidates did not always exhibit the largest fold changes, they occupy functionally central positions within the mitochondrial protein interaction network, and their disruption would be predicted to compromise organelle integrity and function.

**Fig. 3.**
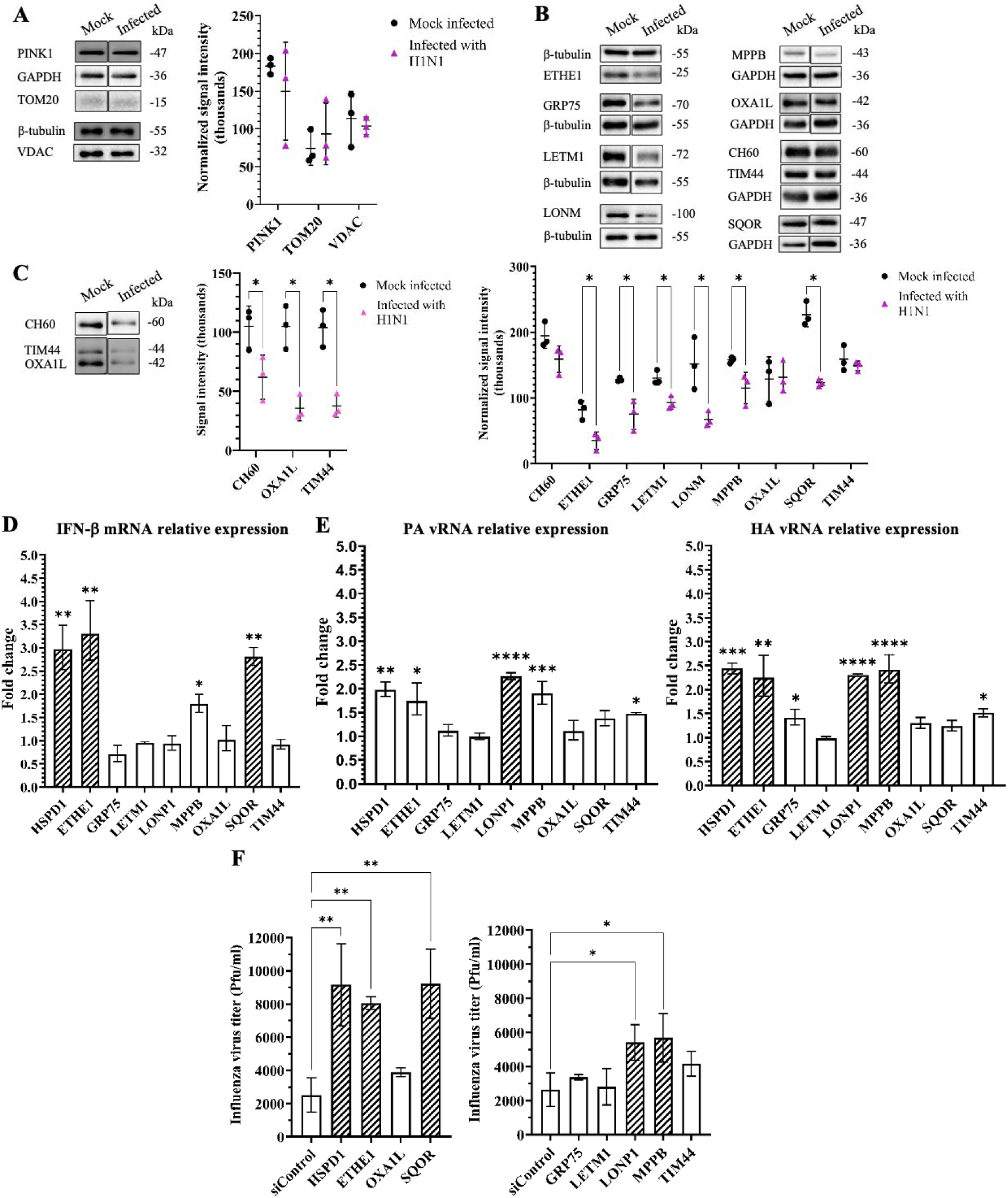
Validation and functional characterization of infection-perturbed mitochondrial proteins. **(A)** Mitochondrial content assessment. To determine whether the observed proteomic changes reflect selective remodeling rather than global organelle loss, levels of the mitophagy marker PINK1 and the mitochondrial mass markers TOM20 and VDAC were assessed by western blot in whole-cell lysates from infected and mock-infected HeLa cells. Values were normalized to GAPDH or β-tubulin. **(B)** Protein-level validation of nine candidates. Abundance of each protein was measured by western blot in whole-cell lysates from infected and mock-infected HeLa cells and normalized to β-tubulin or GAPDH. (**C**) Proximity labeling-specific changes. Abundance of CH60, OXA1L, and TIM44 — which showed stable total cellular levels — was assessed in biotin-phenol-labeled and streptavidin-enriched samples from infected and mock-infected cells. Samples were prepared simultaneously and loaded at equal amounts. In (**A-C**), bars represent normalized means ± s.d., and statistical significance was determined by unpaired two-tailed *t*-test (*p < 0.05). **(D)** Effect on innate immune signaling. HeLa cells were transfected with siRNAs targeting individual candidate genes or a non-targeting control (siControl), infected with influenza A virus, and analyzed for IFN-β mRNA levels by qPCR (normalized to PUM1). (**E**) Effect on viral RNA accumulation. Influenza virus PA and HA vRNA levels were quantified by qPCR in siRNA-transfected and infected HeLa cells (normalized to PUM1). **(F)** Effect on progeny virus production. Infectious virus titers in culture supernatants were determined by plaque assay following infection of siRNA-transfected cells. In (**D-F**), data represent the mean ± SEM of three biological replicates. Patterned bars indicate genes with fold change > 2. Statistical significance was assessed by one-way ANOVA (*p < 0.05, **p < 0.01, ***p < 0.001, ****p < 0.0001).

**Table 1.** Mitochondrial proteins selected for validation and functional characterization.

| Accession | Full protein name | Short protein name | Gene Name | Log2(fold change) |
| --- | --- | --- | --- | --- |
| P10809 | 60 kDa heat shock protein, mitochondrial | CH60 | HSPD1 | -1 |
| O95571 | Persulfide dioxygenase ETHE1, mitochondrial | ETHE1 | ETHE1 | -1.13 |
| P38646 | Stress-70 protein, mitochondrial | GRP75 | GRP75 | -1.36 |
| O95202 | Mitochondrial proton/calcium exchanger protein | LETM1 | LETM1 | -1.25 |
| P36776 | Lon protease homolog, mitochondrial | LONM | LONP1 | -1.25 |
| O75439 | Mitochondrial-processing peptidase subunit beta | MPPB | MPPB | -1.09 |
| Q15070 | Mitochondrial inner membrane protein OXA1L | OXA1L | OXA1L | -1.15 |
| Q9Y6N5 | Sulfide:quinone oxidoreductase, mitochondrial | SQOR | SQOR | -2.06 |
| O43615 | Mitochondrial import inner membrane translocase subunit TIM44 | TIM44 | TIM44 | -1.18 |

We next assessed whether the changes detected by proximity labeling could be confirmed by orthogonal methods. Western blot analysis of total cell lysates from infected and mock-infected cells demonstrated that six of the nine candidates (ETHE1, GRP75, LETM1, LONM, MPPB, and SQOR) were significantly downregulated at the protein level following infection (Fig. 3B). Notably, qPCR analysis revealed no corresponding changes in mRNA expression for five of these six proteins; only LETM1 exhibited a modest but statistically significant decrease in transcript levels (Extended Data Fig. 3). The discordance between protein and mRNA levels for ETHE1, GRP75, LONM, MPPB, and SQOR indicates that their depletion occurs primarily through post-transcriptional mechanism, potentially including accelerated degradation or translational suppression during infection.

The remaining three candidates (CH60, OXA1L, and TIM44) presented a more nuanced picture. Their protein and mRNA levels remained stable in total cell lysates, yet their detection in proximity labeling was significantly reduced in proximity-labeled and enriched mitochondrial fractions from infected cells (Fig. 3C). This discrepancy suggested that infection might alter the subcellular distribution of these proteins, reducing their presence in the mitochondrial matrix without affecting total cellular levels. However, biochemical fractionation of mitochondrial and cytoplasmic compartments revealed no substantial redistribution between conditions (Extended Data Fig. 4).

An alternative interpretation is that infection-induced changes in protein conformation, interaction partners, or complex assembly alter the solvent accessibility of tyrosine residues targeted by biotin-phenoxyl radical, thereby reducing labeling efficiency independently of protein abundance or localization (*27, 28*). Such structural rearrangement would imply that these proteins undergo conformational or interactional remodeling during infection. Accordingly, all nine candidates were carried forward into functional assays.

### Mitochondrial matrix proteins modulate innate immune signal and viral replication

Having established that nine mitochondrial proteins undergo infection-dependent perturbation, we next investigated whether these changes carry functional consequences for the host antiviral response. Mitochondria serve as a central signaling platform in innate immunity: the RIG-I pathway detects cytoplasmic viral RNA and transduces the signal through MAVS, an adaptor anchored to the outer mitochondrial membrane, to induce interferon-β (IFN-β) production (*29, 30*). Whereas the contributions of MAVS and other outer membrane components to antiviral defense have been extensively characterized, the extent to which proteins residing in the mitochondrial matrix and inner membrane influence innate immune signaling remains largely undefined. To address this, we performed siRNA-mediated knockdown of each of the nine candidate genes in HeLa cells, followed by influenza virus infection (Suppl. Fig. 5).

Quantitative PCR analysis revealed that depletion of three candidates (CH60, ETHE1, and SQOR) produced a greater than two-fold increase in IFN-β mRNA levels relative to controls (Fig. 3D). These results indicate that CH60, ETHE1, and SQOR normally function to attenuate interferon signaling, positioning them as negative regulators of the innate immune response operating from within the mitochondrial interior.

Next, we evaluated whether these mitochondrial proteins also influence viral replication directly. Measurement of influenza A virus PA and HA vRNA levels following individual gene knockdowns revealed that depletion of CH60, ETHE1, and MPPB each produced a greater than twofold increase in HA vRNA accumulation relative to siRNA controls (Fig. 3E). The effect was most pronounced for LONM, whose depletion significantly elevated both PA and HA vRNA levels, indicating a broader enhancement of viral genome segment replication in the absence of this protease. To determine whether increased vRNA accumulation translates into greater production of infectious particles, we quantified progeny virus released into the culture supernatant by plaque assay. Knockdown of five proteins (HSPD1, ETHE1, LONP1, MPPB, and SQOR) significantly elevated infectious virus titers compared to controls (Fig. 3F). Notably, this set of proteins overlaps with the three candidates that modulated IFN-β expression: CH60 and ETHE1 influenced both immune signaling and viral replication, whereas LONM and MPPB affected viral output without altering interferon levels, and SQOR elevated both IFN-β expression and progeny virus production. This pattern suggests that the identified mitochondrial proteins restrict influenza virus through at least two distinct mechanisms (immune modulation and direct limitation of viral replication) that are not mutually exclusive (Fig. 4).

**Fig. 4.**
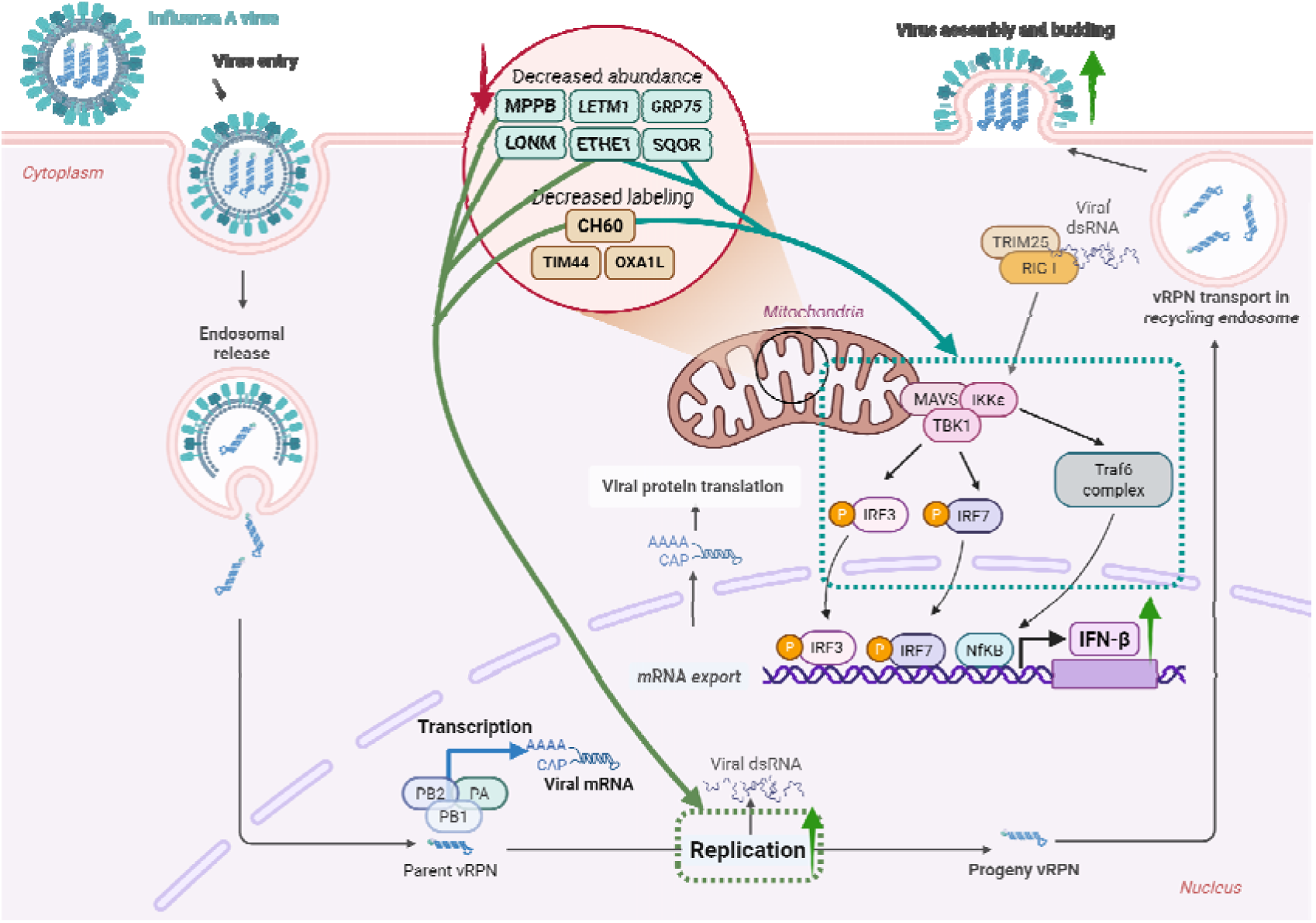
Mitochondrial proteins modulate innate immune response and influenza virus replication. Nine mitochondrial proteins showed reduced cellular abundance or diminished proximity labeling during influenza A virus infection. Functional validation revealed that knockdown of CH60, ETHE1, LONM, MPPB, and SQOR enhanced IFN-β expression and/or vRNA accumulation, ultimately altering the production of progeny virus. Thus, CH60, ETHE1, LONM, MPPB, and SQOR act as mitochondrial restriction factors that modulate host antiviral responses and viral replication.

### Mitochondrial hydrogen sulfide metabolism mediates antiviral restriction

Among the five restriction factors identified above, ETHE1 and SQOR occupy a shared metabolic pathway. Both are essential components of the mitochondrial sulfide oxidation system, in which SQOR catalyzes the initial oxidation of hydrogen sulfide (H□S) to persulfide and ETHE1 subsequently converts the sulfur intermediate to sulfite and thiosulfate.(*31–33*) Their coordinate downregulation during infection and the overlapping phenotypes upon their individual knockdown (e.g. elevated IFN-β expression and increased progeny virus production) suggested that the accumulation of mitochondrial H□S, rather than the loss of each protein per se, might be the proximal effector driving these responses. To test this hypothesis directly, we treated ETHE1- and SQOR-depleted cells with hydroxocobalamin, a cobalamin derivative that functions as a chemical scavenger of H□S, at concentrations of 100 and 200 μM prior to influenza virus infection. Hydroxocobalamin treatment dose-dependently suppressed the elevated viral PA and HA vRNA levels observed upon ETHE1 knockdown, restoring them to near-baseline levels (Fig. 5A, 5B). The elevated IFN-β mRNA expression induced by depletion of either ETHE1 or SQOR was similarly reversed by hydroxocobalamin in a dose-dependent manner (Fig. 5C). Notably, hydroxocobalamin had no significant effect on viral RNA or IFN-β levels in siControl-transfected cells, confirming that the rescue is specific to the H□S-dependent phenotype rather than a general antiviral effect of the compound. These results establish mitochondrial H□S accumulation as the mechanistic link between sulfide pathway disruption and the enhanced viral replication and innate immune activation observed upon ETHE1 and SQOR depletion and identify mitochondrial sulfide metabolism as a previously unrecognized metabolic axis that modulates the host response to influenza A virus infection.

**Fig. 5.**
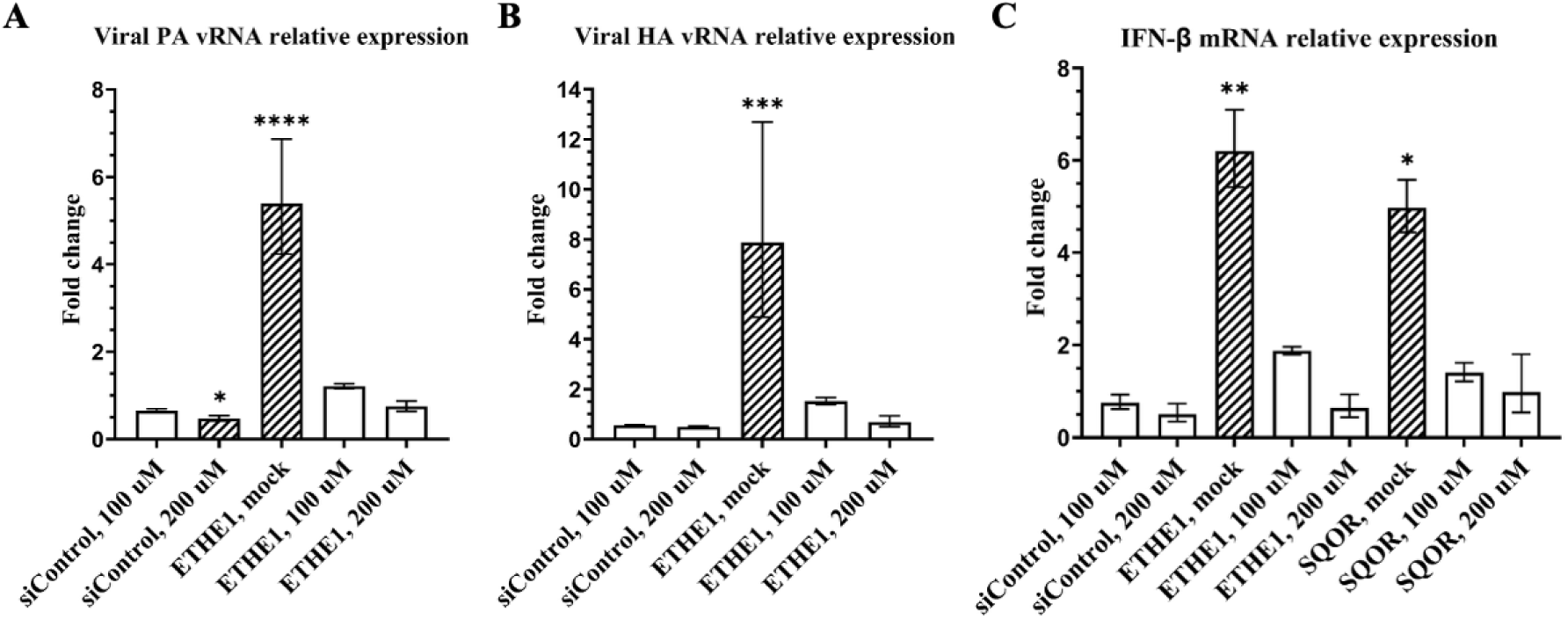
Hydroxocobalamin attenuates elevated viral RNA and IFN-β levels in ETHE1- and SQOR-depleted cells. RT-qPCR analysis of influenza A viral PA and HA vRNAs and IFN-β mRNA in cells transfected with non-targeting control (siControl), ETHE1, or SQOR targeted siRNAs, and treated with increasing concentrations of hydroxocobalamin (0, 100, 200 μM). Expression levels were normalized to siControl mock treated samples and are presented as Log2(Fold change). All data represent the mean ± SEM of three biological replicates. Genes with a fold change > 2 or <0.5 are indicated by patterned bars. Statistical analysis was performed using two-way ANOVA, \**p* < 0.05, \*\**p* < 0.01, \*\*\**p* < 0.001, \*\*\*\**p* < 0.0001.

## Discussion

A central challenge in understanding virus-host interactions is connecting the phenotypic changes visible in infected cells to their underlying molecular origins. Chemosensor-based approaches offer one route to this problem: by targeting specific analytes such as ROS or ER stress markers, they parameterize the response of a defined molecular target with high sensitivity and chemical specificity. However, because each probe interrogates a single predefined analyte, chemosensing strategies inherently constrain the phenotypic search space to what the investigator has chosen to measure. Deep learning-based morphology recognition operates under a fundamentally different logic. Rather than reporting on a preselected molecular event, it learns to extract discriminative features from the full complexity of cellular and organellar shape, capturing phenotypic signatures that may not correspond to any single known biomarker and that may be imperceptible to human observation. In this study, we leveraged this complementary strength through an integrated strategy that pairs image-based deep learning with proximity labeling chemoproteomics. The deep learning component serves as an unbiased phenotyping screen. By training a CNN on organelle-level microscopy images, we identified which subcellular structure undergoes the most pronounced morphological remodeling upon influenza A virus infection, without presupposing the answer. The chemoproteomics component then interrogates the molecular composition of those structures with spatial precision, revealing the specific proteins whose abundance or accessibility is altered within the affected compartment. Together, these two technologies operate at complementary scales, and their sequential application transforms a morphological observation into a mechanistic hypothesis.

Applying this framework to influenza-infected cells, we found that mitochondrial morphology provided the strongest discriminatory signal among the organelles examined, consistently outperforming nuclear, ER, and whole-cell features in classification accuracy. This result directed our molecular investigation toward the mitochondrial matrix, a compartment that has received comparatively little attention in the context of influenza infection, despite the well-characterized involvement of the outer membrane adaptor MAVS in innate antiviral signaling. Proximity labeling of matrix proteome revealed that 99 proteins were coordinately downregulated upon infection, and this remodeling was selective rather than a passive consequence of mitophagy or organelle loss. Meta-analysis also showed that downregulated proteins form functionally coherent modules enriched in mitochondrial organization, respiration, and gene expression - processes whose disruption would be expected to compromise the organelle integrity.

Functional characterization of nine candidates confirmed that the observed chemoproteomic changes carry biological consequences. Five proteins (CH60, ETHE1, LONM, MPPB, and SQOR) emerged as host restriction factors whose depletion enhanced IFN-β expression, viral RNA accumulation or progeny virus production, in several cases affecting multiple axes simultaneously. That their abundance is reduced during infection, while their loss amplifies viral output, supports a model in which influenza virus suppresses mitochondrial restriction factors to create a permissive intracellular environment for its replication. These factors operate through at least two functionally separable mechanisms: i) modulation of innate immune signaling, and ii) direct limitation of viral genome replication. The partial overlap between these two groups (e.g. CH60 and ETHE1 affect both, LONM and MPPB restrict viral RNA, and SQOR primarily modulates IFN-β) suggests a multilayered defense architecture in which the mitochondrial matrix simultaneously communicates with the innate immune apparatus and constrains viral replication through distinct effector arms.

The identification of ETHE1 and SQOR as restriction factors was particularly interesting observation, as both enzymes operate within the same mitochondrial hydrogen sulfide oxidation pathway. They shared elevated IFN-β and increased viral production as phenotypes upon knockdown, together with dose-dependent reversal of these phenotypes by the H_2_S scavenger hydroxocobalamin. This establishes mitochondrial H_2_S accumulation as the proximal metabolic signal linking sulfide pathway disruption to both immune activation and enhanced viral replication. This finding reframes ETHE1 and SQOR not merely as metabolic housekeeping enzymes but as sentinels whose activity levels determine whether mitochondrial H_2_S remains below the threshold that triggers innate immune activation and alters the cellular environment in favor of viral propagation. To our knowledge, this represents the first direct demonstration that mitochondrial sulfide metabolism modulates the host response to influenza virus infection, and it identifies H_2_S as a mitochondria-derived metabolic signal with dual immunomodulatory and proviral functions. On the other hands, the precise mechanism by which CH60, LONM, and MPPB restrict influenza replication remain to be defined. Given their known functions in mitochondrial protein import, folding, and proteolytic quality control, their depletion likely disrupts protein homeostasis within the matrix, potentially triggering both direct effects on viral replication and indirect consequences through mitochondrial stress signaling. Dissecting these individual mechanistic contributions will be an important objective for the future studies.

In summary, this work demonstrates that image-based deep learning and proximity labeling chemoproteomics can be productively combined to trace a path from cellular phenotype to molecular mechanism in the context of viral replication. The approach is inherently generalizable. The deep learning screen identifies the most informative morphological feature without prior biological assumption, and the spatial proteomics component bridges the gap between correlation and causation at molecular resolution. Applied here to influenza A virus, this strategy uncovered a set of mitochondrial matrix proteins that function as host restriction factors and revealed mitochondrial hydrogen sulfide metabolism as a previously unrecognized metabolic axis in antiviral defense. These findings expand the functional landscape of mitochondria in infection biology beyond the outer membrane and into the organelle interior, and they suggest that metabolic pathways operating within the matrix may represent unexplored targets for host-directed antiviral intervention.

## Methods

### Cell culture

HeLa cells, sourced from the Korean Cell Line Bank, were maintained in Dulbecco’s Modified Eagle’s Medium (DMEM) supplemented with 10% (v/v) fetal bovine serum (Gibco) and 1% (v/v) penicillin-streptomycin (Gibco).

### Virus

Konkuk University (Seoul, Republic of Korea) generously provided Influenza A Virus (A/WSN/1933 (H1N1)), 10^7.75^ EID50/ml.

### Plasmid

The plasmid mito-V5-APEX2 (Addgene #72480; http://n2t.net/addgene:72480; RRID: Addgene_72480), gifted by Alice Ting, encodes APEX2 fused at its N-terminus to a 24-amino-acid targeting sequence derived from cytochrome c oxidase subunit 4 isoform 1 (COX4I1). This sequence corresponds to the mitochondrial transit peptide of COX4I1 and directs APEX2 to the mitochondrial matrix. After import, the transit peptide is cleaved by the mitochondrial processing protease, which removes targeting sequences from newly imported precursor proteins. This proteolytic processing generates a characteristic double-band pattern of APEX2 in western blot analysis (Suppl. Fig. 2a).

### Transfection and infection

HeLa cells were seeded into appropriate culture dishes at a density sufficient to achieve 70–80% confluency on the day of transfection. Transfections were performed using the PolyJet transfection reagent (SignaGen Laboratories) following the manufacturer’s standard protocol.

Twenty-four hours post-transfection, the cells were either infected with H1N1 virus (10^6.3^ EID_50_/ml) or mock-infected. The culture medium was removed for infection, and the cells were covered with the viral solution for 45 minutes. Mock-infected samples were treated with PBS without the virus. After incubation, the DMEM High-glucose medium supplemented with 1% antibiotic-antimycotic (Gibco) and 0.05% trypsin (Gibco) was added to the cells. The cells were incubated for 16 hours before proceeding to subsequent experiments.

### Organelle staining for CNN training

HeLa cells were seeded in a black transparent 96-well plate (μGreiner, glass-clear bottom) at a 1 × 10 □cells/well density and cultured in DMEM for 24 hours. To track subcellular organelles, two approaches were utilized: staining with organelle-tracking compounds and transfection with CellLight™ organelle markers. For organelle staining, cells were washed with PBS, infected with the AI virus (10^6^ EID_50_/mL), and incubated for 16 hours. Following infection, cells were treated with 1 μM of various organelle-tracking compounds, including ER-Red BODIPY, ER-Red CMXRos, Lyso-Red DND-99, and ER-Tracker Blue-White (Thermo Fisher Scientific), for 30 minutes. After PBS washes, cells were fixed in 100% methanol for 10 minutes and permeabilized with 0.25% Triton X-100 for another 10 minutes. Alternatively, cells were transfected with CellLight™ organelle markers for 24 hours, including CellLight™ ER-RFP, Mitochondria-RFP, Golgi-RFP, Lysosomes-RFP, and Peroxisome-GFP (Thermo Fisher Scientific). Following transfection, cells were washed with PBS, infected under the same conditions, and subjected to fixation and permeabilization as described above. Nuclei were stained for 15 minutes with Hoechst 33342 (Thermo Fisher Scientific) or for 30 minutes with NUCLEAR-ID Red DNA stain (Enzo, ENZ-52406).

Organelle imaging was performed using a Leica DMi8 microscope (Leica Inc., Germany) with a 40× dry lens. To optimize image acquisition, a mercury lamp (ECL6000, Leica Inc., Germany) provided excitation light at 25% intensity. Images were processed with Leica Application Suite X (LAS X v3.7.0.20979) software. Alternatively, imaging and analysis were conducted using the Operetta High-Content Imaging System (PerkinElmer), with data analysis performed in Harmony 3.1 software (PerkinElmer).

### Dataset preparation and deep learning model training

Images were rescaled to an 8-bit range using min–max normalization. Nuclei were detected using an empirically determined threshold of 25 (8-bit), followed by connected component analysis. Images were cropped around each nucleus, with a maximum size of 150 × 150 pixels, and adjusted in shape as needed to try to avoid including multiple cells, following approaches such as ViresNet (Andriasyan *et al.*^28^). Influenza M protein immunofluorescence images were binarized (threshold > 25), with white pixels marking infected cells, while non-infected cells and background were set to black. To balance classes, datasets were adjusted to a 1:1 ratio of infected and non-infected cells. To improve robustness, data augmentation was applied, including random affine transformations (rotation, translation, scaling, and shear) and horizontal flipping. The final dataset comprised 4,445 mitochondrial, 1,771 endoplasmic reticulum, 10,870 nuclear, and 10,870 bright-field images.

The dataset was split into training (80%), validation (10%), and test (10%) sets. Non-infected cells in the test set were drawn from images without infection. A U-Net-based CNN was implemented in PyTorch, utilizing ∼14.8 million parameters. The model followed an encoder-decoder design with four down-sampling and four up-sampling stages linked by skip connections. Each block consisted of two 3 × 3 convolutions with ReLU activation and batch normalization, and dropout was applied in the decoder layers. The model was trained using the Adam optimizer (learning rate 1 × 10□□), a batch size of 16, and binary cross-entropy loss for up to 10,000 epochs, with early stopping. The final output layer consisted of a 1 × 1 convolution with a sigmoid activation function for classification of infection status.

Model performance was assessed using accuracy, precision, recall, and area under the receiver operating characteristic (ROC) curve. To convert predicted probabilities into binary classifications, an optimal decision threshold was determined from the validation dataset using Youden’s J index. This threshold was then applied to the test set to calculate accuracy, precision, and recall on unseen data. For visualization, model outputs above the threshold were converted into binarized nuclei images representing predicted infected cells, while outputs below the threshold were displayed as blank images.

### Proximity labeling

The medium was removed from the culture dishes, and the cells were incubated in fresh medium supplemented with 1 mM of biotin-phenol (APExBIO). After 1 hour of incubation, the medium was discarded, and the cells were treated with a hydrogen peroxide labeling buffer (1 mM H□O in PBS) for 1 minute. After the treatment, cells were immediately washed three times with freshly prepared quencher solution (10 mM sodium ascorbate (Sigma-Aldrich), 10 mM Trolox (Sigma-Aldrich), 10 mM sodium azide (Sigma-Aldrich) in PBS). Following quenching, the cells were either fixed and permeabilized for immunocytochemistry or harvested by centrifugation, snap frozen, and stored at -80°C for subsequent analyses.

### Immunocytochemistry

The plate wells were blocked with 2.5% bovine serum albumin (BSA, GenDEPOT) for 1 hour at room temperature with gentle shaking. After blocking, cells were incubated for 2 hours with primary antibodies targeting specific proteins, including anti-matrix protein 1 (M1) (1:1000; mouse, Abcam, ab22396), anti-nuclear protein (NP) (1:1000; rabbit, Thermo Fisher Scientific, PA5-32242), anti-V5 tag (1:1000; mouse, Thermo Fisher Scientific, MA5-15253). Unbound antibodies were removed by washing three times with PBS-T (PBS containing 0.1% Tween 20). Cells were then incubated for 1 hour with the appropriate secondary antibodies conjugated to Alexa Fluor dyes (Thermo Fisher Scientific, A11001, A11032, A11012, A11059) at a 1:3000 dilution. Alexa Fluor 488 streptavidin conjugate (Thermo Fisher Scientific, S11223) was used at a 1:2000 dilution for 1 hour to visualize biotin-phenol labeling. After three additional PBS-T washes, nuclei were stained for 15 minutes with either Hoechst 33342 (Thermo Fisher Scientific). Confocal imaging was performed using the LSM800 microscope (ZEISS, Germany) with a C-Apochromat 63×/1.20 W lens and an Airyscan detector. Images were deconvolved using the Airyscan processing method in Zen Blue software. Intensity adjustments and channel alignment were applied when needed.

### Colocalization analysis

Colocalization between Mito-APEX2 and the mitochondrial marker TOM20 was assessed in transfected HeLa cells using confocal fluorescence microscopy images. Only cells that visibly expressed Mito-APEX2 were included in the analysis. Five transfected cells were analyzed from three independent microscopy images. Regions of interest (ROIs) were manually drawn around individual transfected cells using ImageJ (1.53u) to exclude non-expressing cells from the measurements. Colocalization was quantified using the Just Another Colocalization Plugin (JaCoP) in ImageJ. Pearson’s correlation coefficient and Manders’ overlap coefficients (M1 and M2) were calculated to assess the spatial overlap between the red (Mito-APEX2) and green (TOM20) fluorescence channels. Thresholds for calculations were manually adjusted. The resulting coefficients were averaged, and standard deviations were calculated. Data are reported in the graph as mean□±□standard deviation. Individual measurements are reported in Supplementary Table 1.

### Cell lysis

The protocol for sample lysis, enrichment, and mass spectrometry sample preparation was adapted from Kalocsay M.(*34*) Frozen cell pellets were lysed with 8M Urea cell lysis solution. For pellets collected from T150 culture bottles, 0.5 ml of lysis solution was sufficient to dissolve the pellets with gentle pipetting. Next, 0.5 ml of 55% ice-cold trichloroacetic acid (TCA, Sigma-Aldrich) was added to each sample, mixed thoroughly with the lysate, and incubated on ice for 15 minutes. The precipitated proteins were then pelleted by centrifugation at 21,000 x g for 10 minutes at 4°C. The pellets were washed three times with cold acetone (Sigma-Aldrich) and centrifuged under the same conditions. After the final wash, the pellets were dried, and 0.3 ml of lysis solution was added to each pellet. The samples were vortexed for 1 hour at room temperature. Solubilized proteins were centrifuged for 10 minutes at 21,000 x g at 4°C. The clear supernatants were transferred to new tubes. Protein concentrations were determined using the Pierce™ BCA Protein Assay Kit (Thermo Fisher Scientific).

### Immunoblotting

Protein lysates were mixed with Laemmli Sample Buffer (GenDEPOT) boiled at 95°C for 5 minutes, and loaded onto polyacrylamide gels. Proteins were then transferred onto a nitrocellulose membrane (Bio-Rad) at 100V for 1 hour. Transfer efficiency was assessed using Ponceau staining solution. The membranes were subsequently washed once with TBST (Tris-buffered saline with 0.05% Tween 20) and blocked with 3% BSA for 2 hours at room temperature. Following blocking, membranes were incubated overnight at 4°C with primary antibodies. After incubation, the membranes were washed 3 times with TBST for 10 minutes. Then, membranes were incubated with secondary antibodies conjugated to either Alexa Fluor dyes or HRP (Thermo Fisher Scientific, A11001, A32731, A32728, A21245, G21234; Jackson ImmunoResearch, 115-035-166) at 1:5000 dilution for 1 h. Alexa Fluor 488 streptavidin conjugate (Thermo Fisher Scientific, S11223) was used at a 1:3000 dilution for 30 minutes to visualize biotin-phenol labeling. After incubation, membranes were washed three times. HRP signals were detected using Pierce ECL Western Blotting Substrate (Thermo Fisher Scientific), and protein bands were visualized using the ChemiDoc XRS+ system (Bio-Rad) with Image Lab software v6.1.0. Protein band intensity was analyzed using Image Lab or ImageJ v1.53u software. Primary antibodies used:

anti-GAPDH (1:2000; rabbit, Cell Signaling Technology, 2118)

anti-beta Tubulin (1:2000; rabbit, Abcam, ab15568)

anti-beta Tubulin (1:2000; mouse, Thermo Fisher Scientific, MA5-16308)

anti-V5 tag (1:1000; mouse, Thermo Fisher Scientific, MA5-15253)

anti-CH60 (1:6000; rabbit, Proteintech, 15282-1-AP)

anti-ETHE1 (1:1000; rabbit, Proteintech, 27786-1-AP)

anti-GRP75 (1:500; mouse, Thermo Fisher Scientific, MA3-028)

anti-LETM1 (1:1000; rabbit, Proteintech, 16024-1-AP)

anti-LONM (1:1000; rabbit, Proteintech, 15440-1-AP)

anti-OXA1L (1:2000; rabbit, Proteintech, 21055-1-AP)

anti-MPPB (1:1000; rabbit, Proteintech, 16064-1-AP)

anti-SQOR (1:1000; rabbit, Proteintech, 17256-1-AP)

anti-TIM44 (1:1000; rabbit, Abcam, ab244466)

anti-TOMM20 (1:500; mouse, Abcam, ab56783)

anti-VDAC (1:1000; rabbit, Cell Signaling Technology, 4661).

### Streptavidin pulldown

A total of 1.5 mg of protein lysates were volume-equalized and treated with 20 μM iodoacetamide for 25 minutes at room temperature with gentle shaking. Then, 50 μM dithiothreitol (DTT) was added, and incubation continued for 15 minutes. The samples were diluted to achieve a final urea concentration of 2 M. Streptavidin-based enrichment was performed using 300 μL of Pierce streptavidin magnetic beads (Thermo Fisher Scientific). The beads were washed twice with equilibration buffer (2 M urea, 0.25% SDS in sodium phosphate buffer) before adding the protein samples. The samples were incubated with the beads overnight at 4°C with gentle rotation. After incubation, the beads were sequentially washed: first wash with 2 M urea, 0.25% SDS buffer, second wash with 2 M urea buffer, third wash with 1 M urea buffer. To assess biotinylated protein capture, 5% of the beads were separated and eluted with 1× Laemmli Sample Buffer, followed by boiling at 95°C for 5 minutes. The remaining beads were used for on-bead trypsin digestion.

### Mass spectrometry sample preparation

Streptavidin beads were resuspended in 200 μL of digestion buffer (1 M urea, 200 mM HEPES pH 8.5, 4% (v/v) acetonitrile) supplemented with 2 μg of trypsin (Promega). Samples were incubated at 37°C for 3 hours with shaking. After digestion, the supernatant was transferred to new tubes, and the beads were washed twice with 100 μL of digestion buffer. The washes were pooled with the initial digestion supernatant. An additional 1.5 μg of trypsin was added, and digestion was continued for 16 hours at room temperature with shaking.

Following overnight digestion, peptides were acidified with formic acid (Thermo Fisher Scientific) to pH 2.5 and desalted using Sep-Pak C18 cartridges (Waters). The cartridges were conditioned with 1 mL of 100% acetonitrile and 1 mL of 0.1% formic acid. Peptides were loaded, washed with 1 mL of 0.1% formic acid, and eluted with 400 μL of 80% acetonitrile, 0.1% formic acid solution. The eluates were vacuum-dried and reconstituted in 50 μL of 200 mM HEPES pH 8.5.

For isobaric labeling, each sample was labeled with an individual TMT reagent (Thermo Fisher Scientific, TMT10plex™ Isobaric Label Reagents and Kit, 90110) for 1 hour at room temperature. The reaction was quenched by adding 5 μL of 10% (v/v) hydroxylamine and incubating for 15 minutes at room temperature with shaking. The labeled samples were then pooled, acidified with formic acid to pH 2.5, and vacuum-dried.

The samples were processed using the Pierce High pH Reversed-Phase Peptide Fractionation Kit (Thermo Fisher Scientific) according to the manufacturer’s protocol. Peptides were eluted in fractions containing increasing acetonitrile concentrations (10%, 12.5%, 15%, 17.5%, 20%, 25%, 50%, and 80%). The fractions were vacuum-dried and stored at -80°C until further analysis.

### Mass spectrometry

Dried peptides were resuspended in 10 μL of 0.1% formic acid. Chromatographic separation was performed using a 50-cm C18 analytical column (Thermo Fisher Scientific) and a 2-cm trap column (Thermo Fisher Scientific), with the analytical column maintained at 50°C. Peptides were eluted using a 180-minute gradient ranging from 5% to 80% acetonitrile in 0.1% formic acid. LC-MS/MS/MS data were acquired using an UltiMate™ 3000 RSLCnano System (Thermo Fisher Scientific) coupled to an Orbitrap Eclipse Tribrid mass spectrometer (Thermo Fisher Scientific) and controlled by Xcalibur software v4.1. Full-scan MS spectra were collected in the m/z range of 400–2,000 at a resolution of 120,000 in the FT-Orbitrap. The twenty most intense ions were selected for HCD-MS2 fragmentation, followed by the selection of the ten most abundant fragment ions for HCD-MS3 analysis. HCD-MS3 acquisition was employed to minimize TMT ratio compression, as it reduces interference from co-isolated peptides.

MS3 data were searched against the Uniprot Homo sapiens (Human) and Influenza A virus (A/WSN/1933(H1N1)) database (2023.02.22) using Proteome Discoverer software v2.4. Search parameters included TMT-plex (K) and carbamidomethylation (C) as fixed modifications. Oxidation (M) and acetyl (N-terminus) were set as variable modifications. Precursor and fragment mass tolerances were set to 10 ppm and 0.9 Da, respectively. Peptide and site false discovery rates were controlled at 1% using Percolator. Total Ion Current normalization was performed to account for variation in peptide amounts.

### Receiver operating characteristic (ROC) curve analysis

The data analysis was adapted from Cho K. F. *et al.* (*35*) Briefly, TMT ratios were calculated by dividing the TMT signal from the experimental sample by the signal from the negative-control sample to identify biotinylated proteins. The ratios were then ranked and compared to curated lists of true positive and false positive proteins, generated based on prior annotation and evidence from the literature. The ROC curve was plotted to determine the optimal TMT ratio cutoff, which maximized the true positive rate while minimizing the false positive rate. Proteins above the cutoff were considered biotinylated, while those below were considered nonspecific binders.

### Gene knockdown

HeLa cells were seeded in 6-well plates 24 hours before transfection. Once the cells reached approximately 75% confluency, they were transfected with targeted or random siRNA using siTrans 2.0 transfection reagent (OriGene) according to the manufacturer’s protocol. The final siRNA concentration was 10 nM per well. The medium was changed 18 hours after transfection. The cells were collected after 48 hours to analyse for knockdown efficiency or used for further experiments.

siRNA used:

Hsp60 (HSPD1) Human siRNA Oligo Duplex (OriGene, locus ID 3329)

AccuTarget Predesigned Human ETHE1 siRNA (Bioneer, locus ID 23474)

AccuTarget Predesigned Human GRP75 siRNA (Bioneer, locus ID 3313)

LETM1 Human siRNA Oligo Duplex (OriGene, locus ID 3954)

AccuTarget Predesigned Human LONP1 siRNA (Bioneer, locus ID 9361)

OXA1L Human siRNA Oligo Duplex (OriGene, locus ID 5018)

AccuTarget Predesigned Human MPPB siRNA (Bioneer, locus ID 9512)

AccuTarget Predesigned Human TIM44 siRNA (Bioneer, locus ID 10469)

AccuTarget Predesigned Human SQOR siRNA (Bioneer, locus ID 58472)

Non-targeting scramble siRNA (siControl) was purchased from OriGene.

### Real-time quantitative PCR

After gene knockdown, HeLa cells were infected with H1N1 virus (10^6.3^ EID_50_/ml) for 16 hours in triplicate. After infection total RNA was isolated from the cells using the RNeasy Plus kit (QIAGEN) following the manufacturer’s protocol. RNA concentration was measured using a spectrophotometer, and the A260/A280 ratio was assessed to confirm sample purity. Reverse transcription was carried out using the TOPscript cDNA Synthesis Kit (Enzynomics). Each reaction used 1 μg of total RNA, and cDNA synthesis was performed on a T100 Thermal Cycler (Bio-Rad).

Quantitative PCR reactions were prepared using TOPreal™ SYBR Green qPCR PreMIX (Enzynomics). The qPCR cycling protocol included an initial denaturation at 95°C for 10 minutes, followed by 40 denaturation, annealing, and extension cycles, with an annealing temperature of 60°C. Amplification was performed on a QuantStudio 3 Real-Time PCR System (Thermo Fisher Scientific).

Amplification curves were analyzed using Design & Analysis Software v2.7.0 (Thermo Fisher Scientific) with the automatic threshold method applied. Relative gene expression levels were calculated using the ΔΔCt method, normalizing target gene Ct values to Pumilio RNA Binding Family Member 1 (PUM1) as the reference gene. Mean, standard deviation, standard error of the mean, and statistical tests were calculated using ΔCt values. The values were transformed to a linear scale and plotted on graphs.

qPCR oligonucleotides for the knockdown of targeted genes were purchased from Bioneer. Additional primers were purchased from Macrogen, including those targeting PUM1 (5’-TGCGGGAGATTGCTGGACAT-3’, 5’-GTGTGGCACGCTCCAGTTTC-3’), Influenza HA segment (5’-CGAAGACAGACACAACGGGA-3’, 5’-GTTTGGTGTTTCTACAATGTAGGACC-3’), Influenza PA segment (5’-GGATTCGAACCGAACGGCTAC-3’, 5’-TTTGGACCGCTGAGAACAGG-3’), Interferon beta 1 (5’-AGCTGCAGCAGTTCCAGAAG-3’, 5’-AGTCTCATTCCAGCCAGTGC-3’).

### Plaque assay

Following gene knockdown, HeLa cells were infected with the H1N1 virus (10^6.3^ EID □ □/mL) for 16 hours in triplicate. Culture media were collected and concentrated using Amicon Ultra-4 100K Centrifugal Filters (Merck Millipore) by centrifugation at 7,500 × g for 15 minutes. The final sample volumes were equalized with PBS.

MDCK cells were seeded in 6-well plates at a density of 0.4 × 10 cells per well. After 24 hours, viral samples were mixed 1:1 with virus growth medium (DMEM supplemented with 0.15% (v/v) bovine serum albumin, 25 mM HEPES, and 1% (v/v) penicillin-streptomycin) and subjected to a 2-fold serial dilution. Cells were infected with 400 μL of each viral dilution or mock-infected with PBS in duplicate and incubated at 37°C for 45 minutes. Following incubation, each well received a 3 mL overlay, prepared by mixing equal volumes of 2X DMEM (26.8 g DMEM powder, 7.4 g sodium bicarbonate, 0.375% (v/v) BSA, 50 mM (v/v) HEPES, 10 mM (v/v) GluMAX, 1% (v/v) penicillin-streptomycin) and 2.4% (w/v) cellulose (Sigma-Aldrich), supplemented with 1.5 μg/mL TPCK-trypsin (Thermo Fisher Scientific). Plates were incubated at 37°C for 72 hours.

After incubation, wells were fixed with 3 mL of 10% formaldehyde for 1 hour, aspirated, washed with PBS, and stained with 1% (w/v) crystal violet (Sigma-Aldrich) for 10 minutes. Wells were then washed with water and allowed to dry. Plaques were manually counted. Viral titers were calculated by multiplying the average number of plaques by the reciprocal dilution factor and the inoculum volume.

### Statistics

Statistical analysis was performed using GraphPad Prism software v10.4.0, with results expressed as the mean value ± standard deviation or standard error of the mean. A one-way analysis of variance (ANOVA) was employed to determine statistical significance, with a threshold of *p* < 0.05 indicating statistical significance. For comparisons involving multiple groups, Dunnett’s test was used. Pairwise comparisons were evaluated using an unpaired t-test, with a threshold of *p* < 0.05 indicating statistical significance. Welch’s correction was applied in case of unequal variance between groups.

## Supporting information

Supplementary Images

## Data availability

The mass spectrometry data were deposited to MassIVE (dataset MSV000098251, password: Gcnth79DAoSUz1X2).

## Code availability

All scripts and detailed hyperparameter settings are available at Github [https://github.com/Imfinethankyou1/Influenza-A-Virus-Infection-U-Net.git].

## Acknowledgments

This work was supported by the Ministry of Science and ICT through the National Research Foundation of Korea (RS-2025-00523442, RS-2025-16652968, RS-2023-00217701, RS-2025-02305395, RS-2024-00403511) and a National Research Council of Science and Technology (NST) grant (CAP23011-400).

## Author contributions

O.S., M.M.H., K.T.H. performed the experiments; H.K. built and tested CNN model; O.S., Y.K. analyzed the data; W.Y.K., J.-S.L. designed the study; O.S. wrote the original draft of the manuscript; O.S., Y.K. and J.-S.L. wrote and edited the manuscript.

---

**Extended Data Fig. 1.**
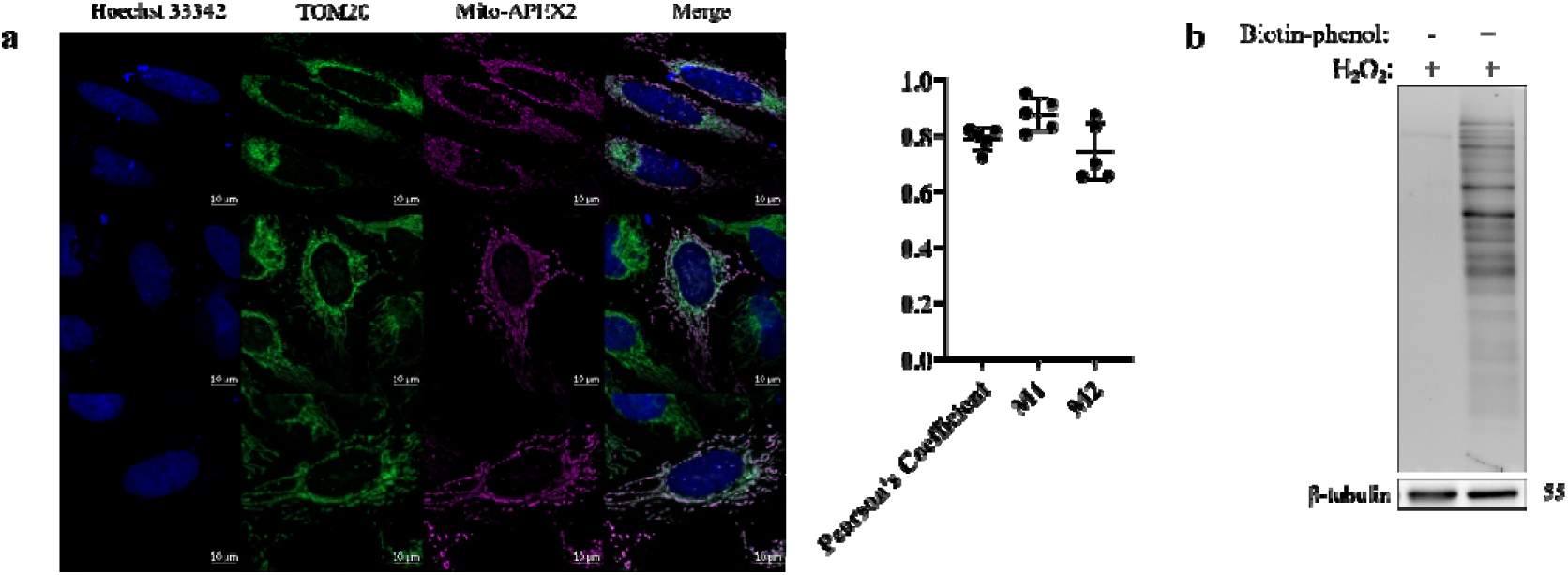
Characterization of APEX2 construct. **a,** Confocal fluorescence imaging of HeLa cells expressing APEX2 targeted to mitochondria. APEX2 expression was visualized using an anti-V5 tag antibody, TOM20 was labeled with anti-TOM20 antibody. Colocalization was analyzed in five transfected HeLa cells using the JaCoP plugin in ImageJ. Regions of interest (ROIs) were manually defined to include only cells expressing Mito-APEX2. Pearson’s correlation coefficient and Manders’ overlap coefficients (M1: proportion of Mito-APEX2 signal overlapping with TOM20; M2: proportion of TOM20 signal overlapping with Mito-APEX2) were calculated using manually defined thresholds. In the graph, the bars represent means ± s.d. of five measurements for each coefficient. **b,** Characterization of APEX2 labeling activity. HeLa cells expressing Mito-APEX2 were treated with biotin-phenol or mock-treated (medium without compound) for 1 hour, followed by incubation with 1 mM HDOD for 1 minute. Whole-cell lysates were analyzed by Alexa Fluor 488 streptavidin blotting. Equal protein loading was confirmed by β-tubulin detection.

**Extended Data Fig. 2.**
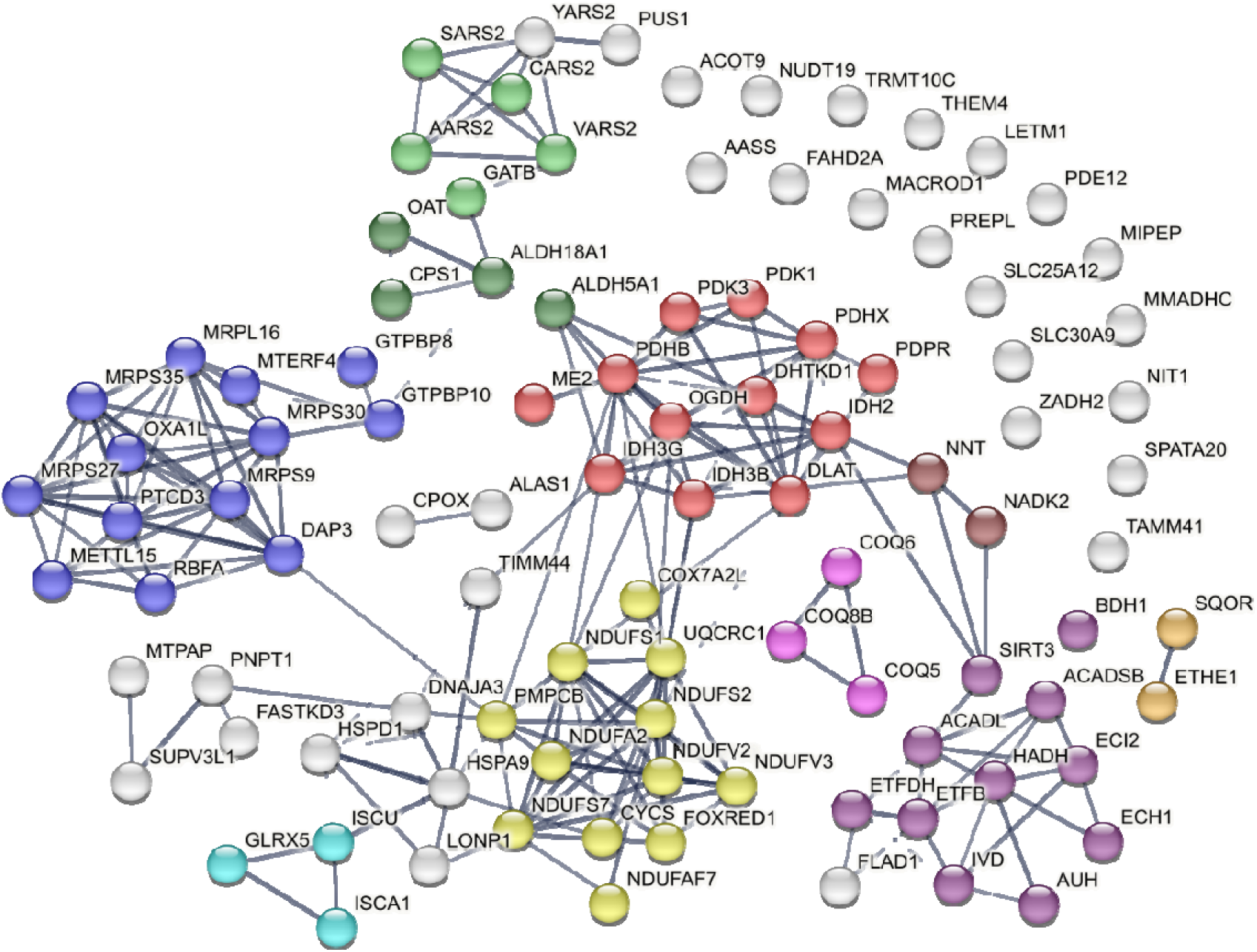
STRING network analysis. The node colors represent local network clusters (STRING). Ten non-overlapping clusters were chosen for display: citrate cycle and pyruvate metabolism (FDR = 3.25e-12, red), organellar ribosome and regulation of mitochondrial gene expression (FDR = 1.64e-10, blue), fatty acid beta-oxidation and AMP-binding (FDR = 5.91e-09, violet), respiratory electron transport and ATP synthesis (FDR = 3.32e-08, yellow), tRNA aminoacylation (FDR = 4.5e-04, light green), ubiquinone biosynthesis (FDR = 4.3e-03, pink), glutamine family amino acid biosynthesis (FDR = 6.2e-03, dark green), iron-sulfur cluster assembly (FDR = 3.95e-02, cyan), NAD kinase and NAD/GMP synthase (FDR = 3.97e-02, brown), hydrogen sulfide metabolic process (FDR = 3.97e-02, dark yellow).

**Extended Data Fig. 3.**
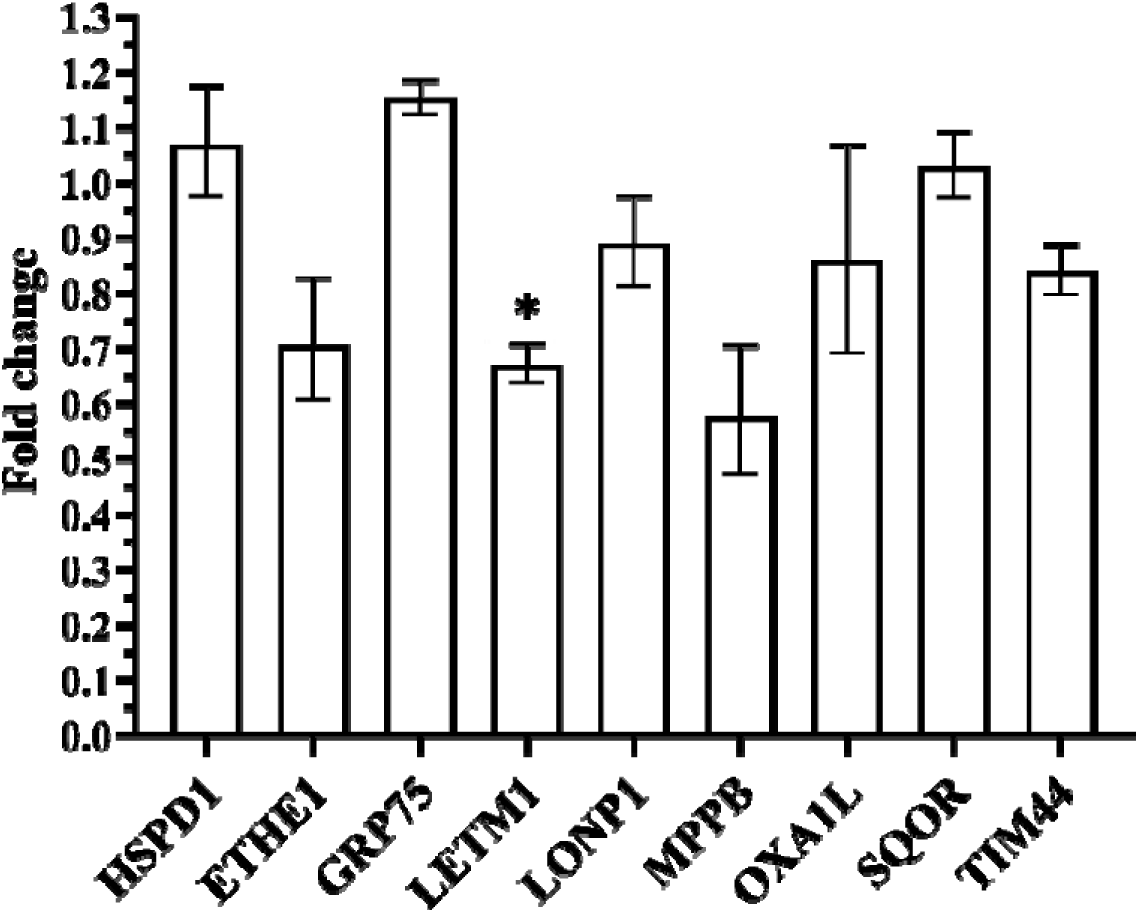
Fold change in mRNA levels of target proteins in infected versus mock-infected cells. Total RNA was extracted from infected and mock-infected HeLa cells, and relative mRNA levels were quantified by RT-qPCR using PUM1 as the reference gene. Expression levels in mock-infected samples were set to 1, and infected sample values are shown as fold change relative to mock. Data represent mean ± s.e.m. from three biological replicates. Statistical significance was determined using multiple t-tests, *p < 0.05.

**Extended Data Fig. 4.**
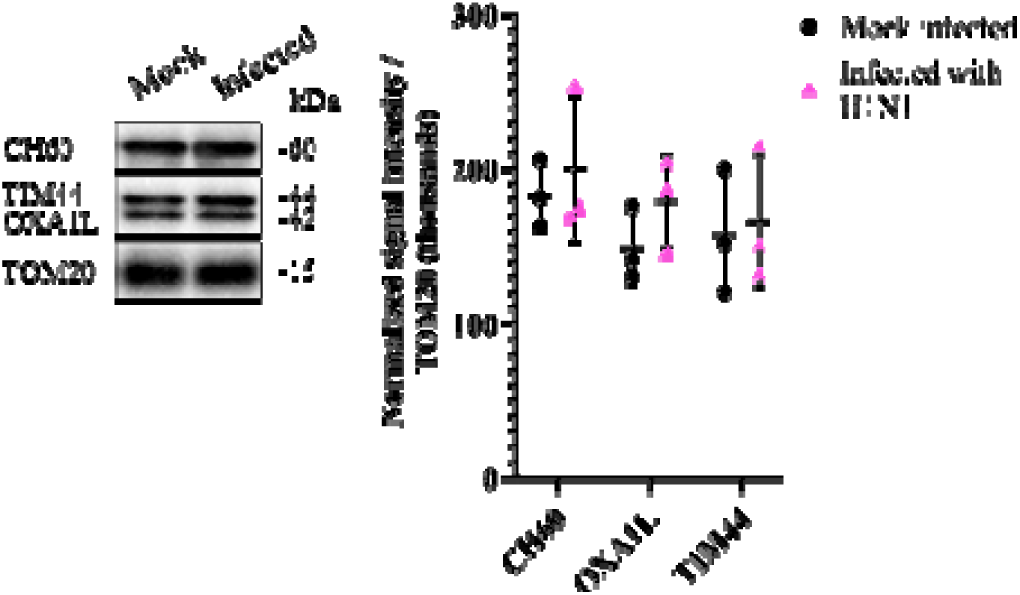
Mass spectrometry validation. Abundance levels of CH60, OXA1L and TIM44 proteins were calculated in mitochondrial fractions. The values were normalized with TOM20 levels. The bars on all graphs represent the normalized means ± s.d. of triplicate samples. Statistical significance was calculated using a parametric unpaired t-test, *p < 0.05.

