## Supplementary Images for "Cell morphology deep learning guides discovery of mitochondrial host defense against influenza"

Supplementary Figures 1-5

Supplementary Table 1-2

Supplementary Tables 3-4 Legends

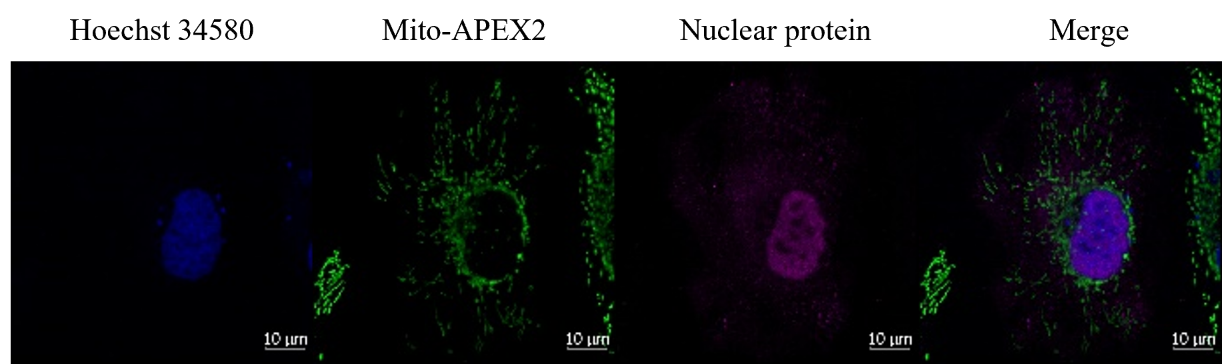

**Supplementary Figure 1.** Confocal fluorescence imaging of infected HeLa cells expressing APEX2 targeted to mitochondria. APEX2 expression was visualized using an anti-V5 tag antibody, while infected cells were identified with an anti-nuclear protein antibody.

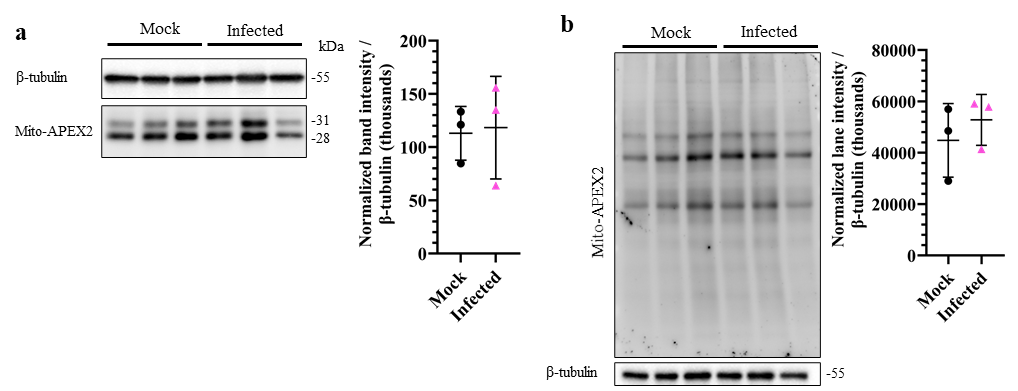

**Supplementary Figure 2.** **a,** Level of Mito-APEX2 expression in infected and mock-infected cells. Expressed APEX2 was detected in cell lysate samples using anti-V5 tag antibody. The values were normalized with β-tubulin levels. **b,** Level of Mito-APEX2 labeling activity in infected and mock-infected cells. HeLa cells expressing APEX2 were either infected or mock-infected for 24 hours before biotin-phenol labeling. The labeling intensities were analyzed by Alexa Fluor 488 streptavidin blotting. The bars on the graphs represent the normalized means ± s.d. of triplicate samples. Statistical significance was calculated using a parametric unpaired t-test.

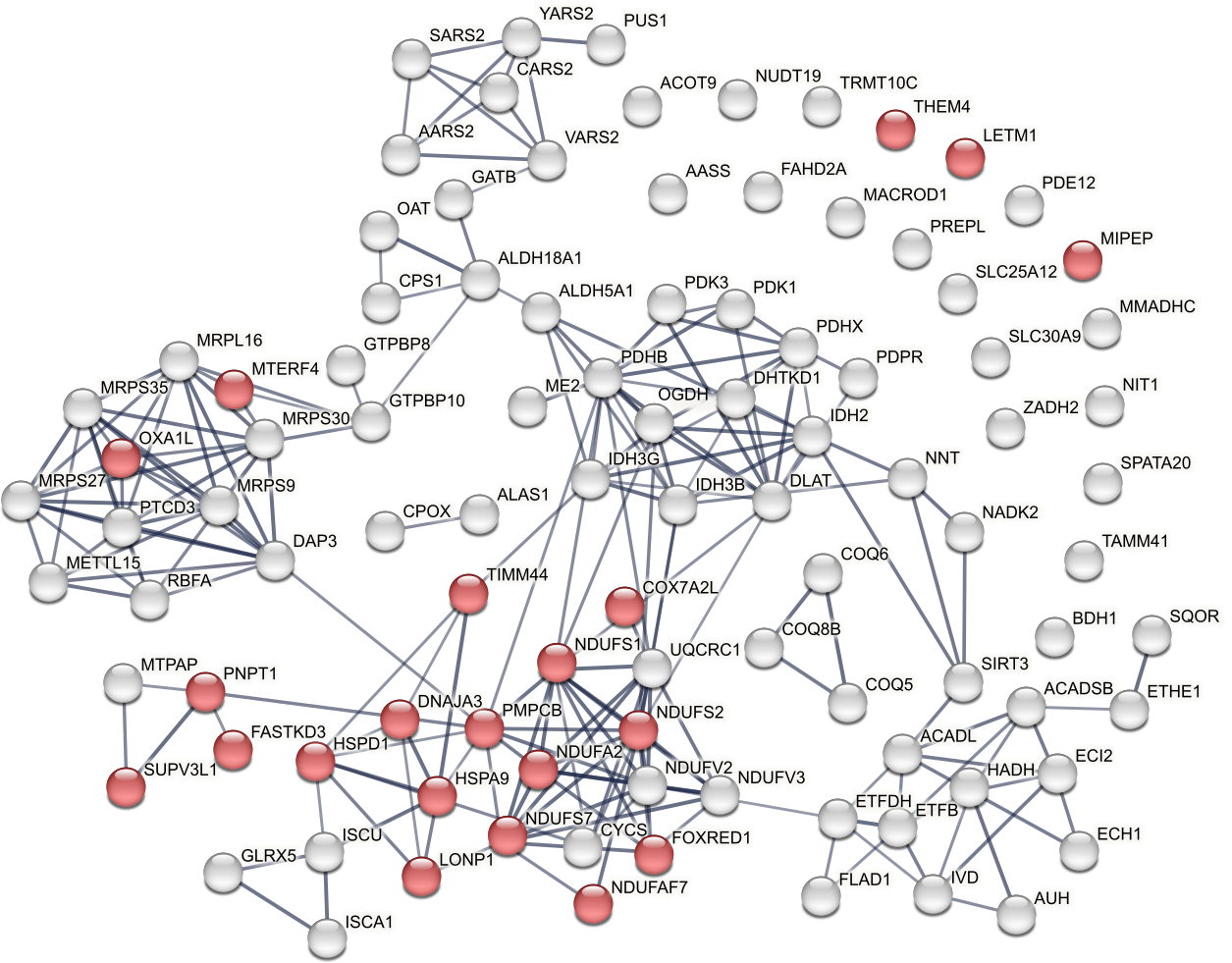

**Supplementary Figure 3.** STRING network analysis. The edges indicate functional and physical protein associations, and the line thickness indicates the strength of the data support. The minimum required interaction score was set at 0.700. Red node color indicates proteins involved in ‘Mitochondrion organization’ (GO:0007005) category (FDR = 1.36e-11).

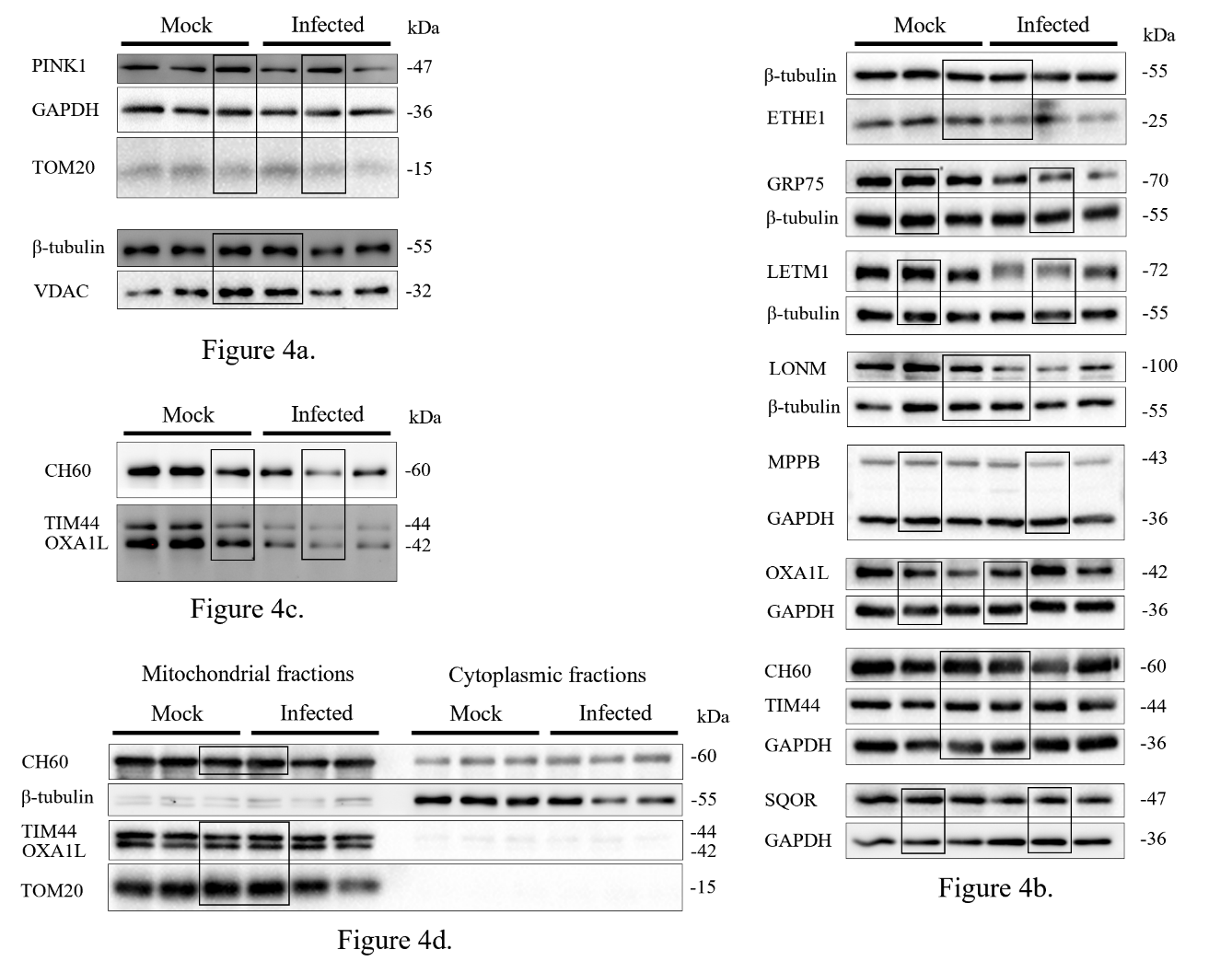

**Supplementary Figure 4.** Original Western blot images used in the preparation of Figure 4, with the relevant sections indicated by a black box.

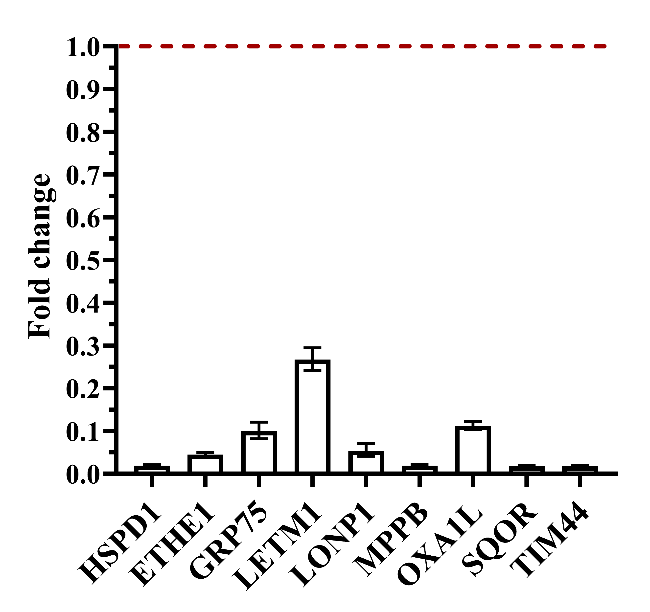

**Supplementary Figure 5.** HeLa cells were transfected with the indicated siRNAs. siRNA knockdown efficiencies were analyzed with RT-qPCR 48 hours after transfection. Expression was normalized to the reference gene (PUM1). Control expression levels were set to 1, and knockdown values are shown relative to control. Error bars represent means ± SEM, biological triplicate.

| **Input** | **AUROC** | **Recall** | **Precision** | **Accuracy** |
| --- | --- | --- | --- | --- |
| Bright field | 0.723 | 0.464 | 0.734 | 0.674 |
| Nuclei | 0.789 | 0.783 | 0.634 | 0.691 |
| Mitochondria | 0.854 | 0.714 | 0.849 | 0.797 |
| Endoplasmic reticulum | 0.671 | 0.753 | 0.534 | 0.596 |

**Supplementary Table 1.** Performance metrics of the deep learning model evaluated separately for each image type used as training input.

| **Cell** | **Pearson's Coefficient** | **M1** | **M2** |
| --- | --- | --- | --- |
| 1 | 0.782 | 0.830 | 0.876 |
| 2 | 0.823 | 0.808 | 0.702 |
| 3 | 0.723 | 0.881 | 0.659 |
| 4 | 0.803 | 0.914 | 0.657 |
| 5 | 0.818 | 0.952 | 0.833 |
| Mean | 0.790 | 0.877 | 0.745 |
| SD | 0.041 | 0.059 | 0.102 |
| Lower 95% CI of mean | 0.739 | 0.804 | 0.618 |
| Upper 95% CI of mean | 0.840 | 0.950 | 0.872 |

**Supplementary Table 2.** Colocalization analysis of Mito-APEX2 and TOM20 in transfected HeLa cells. Colocalization was assessed in five individually analyzed transfected HeLa cells using the JaCoP plugin in ImageJ. Regions of interest (ROIs) were manually drawn around Mito-APEX2-expressing cells. Pearson’s correlation coefficient and Manders’ overlap coefficients (M1: fraction of Mito-APEX2 overlapping with TOM20; M2: fraction of TOM20 overlapping with Mito-APEX2) were calculated using a manually set threshold.

**Supplementary Table Legends**

**Supplementary Table 3. Mass spectrometry results, mitochondria.**

Tab 1: "Proteins" – Lists all identified proteins with at least two unique peptides.

Tab 2: "APEX2 vs Control" – Contains proteins significantly enriched in APEX2-expressing samples compared to controls (Log₂(Fold change) > 1, p < 0.05).

Tab 3: "GO" – Filters the list from Tab 2 to include only proteins annotated with the mitochondrial cellular component, based on Gene Ontology (GO) analysis.

Tab 4: "APEX2 Inf vs APEX2" – Lists proteins with significantly reduced abundance in influenza-infected versus mock-infected APEX2 samples (Log₂(Fold change) ≤ –1, p < 0.05).

Tab 5: "ROC Method" – Shows calculations used in the Receiver Operating Characteristic (ROC) curve analysis.

Tab 6: "APEX2 vs Control ROC" – Includes proteins that passed the ROC-based cutoff for specific mitochondrial labeling.

Tab 7: "Comparison with Mito True Pos" – Compares two identified mitochondrial protein lists (from Tabs 2 and 6) with a reference set of known mitochondrial proteins (“Mitochondrial true positive”). This data supports the Venn diagram shown in Fig. 3a.

Tab 8: "Volcano Plot" – Contains the data used to generate the volcano plot displayed in Fig. 3b.

Tab 9: "APEX2 Inf vs APEX2 Expanded" – Lists proteins from the expanded ROC-based mitochondrial dataset that are significantly downregulated in infected samples (Log₂(Fold change) ≤ –1, p < 0.05).

Tab 10: "Target Proteins" – Provides functional annotations for proteins in the previous tab. The nine proteins selected for further validation are highlighted in green.

**Supplementary Table 4. STRING interactions analysis results.**

Tab 1: "Proteins" – Lists mitochondrial proteins along with their fold change values that were used as input for STRING network analysis.

Tab 2: "STRING Interactions" – Contains all protein–protein interactions identified by STRING for the input dataset.

Tab 3: "GO Biological Process" – Includes all enriched Gene Ontology (GO) biological process terms associated with the STRING network. The term “Mitochondrion organization” is highlighted in orange.

Tab 4: "GO Molecular Function" – Lists all GO molecular function terms significantly enriched within the STRING network.

Tab 5: "GO Cellular Component" – Contains enriched GO cellular component terms identified in the STRING analysis.

Tab 6: "Local Network Clusters (STRING)" – Shows all clusters identified by the STRING network clustering algorithm. The clusters shown in Extended Data Fig. 2 are highlighted in their respective colors.
